# Standard Procedures for Native CZE-MS of Proteins and Protein Complexes up to 800 kDa

**DOI:** 10.1101/2020.11.02.351247

**Authors:** Kevin Jooss, John P. McGee, Rafael D. Melani, Neil L. Kelleher

## Abstract

Native mass spectrometry (nMS) is a rapidly growing method for the characterization of large proteins and protein complexes, preserving “native” non-covalent inter- and intramolecular interactions. Direct infusion of purified analytes into a mass spectrometer represents the standard approach for conducting nMS experiments. Alternatively, CZE can be performed under native conditions, providing high separation performance while consuming trace amounts of sample material. Here, we provide standard operating procedures for acquiring high quality data using CZE in native mode coupled online to various Orbitrap mass spectrometers via a commercial sheathless interface, covering a wide range of analytes from 30 – 800 kDa. Using a standard protein mix, the influence of various CZE method parameters were evaluated, such as BGE/conductive liquid composition and separation voltage. Additionally, a universal approach for the optimization of fragmentation settings in the context of protein subunit and metalloenzyme characterization is discussed in detail for model analytes. A short section is dedicated to troubleshooting of the nCZE-MS setup. This study is aimed to help normalize nCZE-MS practices to enhance the CE community and provide a resource for production of reproducible and high-quality data.

## 1 Introduction

Over the last two decades, native mass spectrometry (nMS) has become a powerful analytical method to analyze large proteins and protein complexes [1–3]. Volatile, aqueous buffer systems close to neutral pH are used to preserve endogenous (“native”) characteristics of proteins and their non-covalent interactions [4,5]. In this way, the composition of biomolecular structures can be studied in detail, leading to a better understanding of their topology, stoichiometry, and biological functions [6,7]. There are distinct differences in the nature of protein mass spectra obtained under native conditions compared to traditional denaturing conditions [8]. Fewer protonation events take place during native electrospray ionization, and consequently, lower and fewer charge states for proteins are observed [9]. This characteristic can lead to increased sensitivity, especially for large proteins and their complexes. The mass-to-charge ratio (*m*/*z*) spacing between lower charge states of a native protein distribution is increased compared to the higher charge states of denatured counterparts, which can be beneficial analyzing proteins with similar mass or structure (e.g. proteoforms).

Native top-down mass spectrometry (nTDMS) enables the characterization of protein complexes and their interaction sites, which can also provide insight into interactions with metals, ligands, substrates, reaction products and drug fragments [10–13]. Furthermore, protein complexes can consist of different proteoforms arising from posttranslational modifications, single nucleotide polymorphisms, and RNA splicing variants and thus, can be considered as multi-proteoform complexes (MPCs) [14,15]. The process of ejecting subunits of a protein complex followed by targeted fragmentation is often referred to as “complex-down” and will be considered part of the nTDMS tool box [16]. Developments in mass analyzer technologies [17,18], including time-of-flight [19], ion cyclotron resonance [20], and Orbitrap [21], have broadened the field of application for nMS to protein assemblies with a wide range in size, shape, complexity, and throughput. In addition, various fragmentation techniques and their combination improved the capabilities of subunit ejection and subsequent fragmentation in nTDMS [22–26]. Despite all these advancements in technology, ion suppression and signal superposition cannot be completely avoided in a direct infusion nMS experiment, especially for complex samples. Thus, upfront separation under native conditions is beneficial and highly desired.

There are many analytical separation techniques that can be compatible with native separation. “Native-friendly” chromatographically based separation approaches include size exclusion chromatography [27], hydrophobic interaction chromatography [28], ion exchange chromatography [29], and affinity chromatography [30]. In addition, electromigrative separation techniques such as capillary isoelectric focusing [31], native Gel-Eluted Liquid Fraction Entrapment Electrophoresis (GELFrEE) [32], flow field-flow fractionation [33], and CZE [34,35] have been applied for native protein analysis. In this context, three major prerequisites must be fulfilled to achieve efficient native separation coupled to an MS instrument: (i) the buffer/electrolytes must preserve native features of the protein or protein assembly; (ii) the composition of the spray solution must be compatible with electrospray ionization to avoid ion suppression; and (iii) the separation performance and resolution need to be sufficient to separate targeted analytes. CZE fulfils all three aspects and can provide high-resolution even under native conditions [35] while consuming a few nanoliters per injection and exhibiting low carry-over [36]. In CZE, separation takes place based on differences in electrophoretic mobilities of ions in liquid phase, which is dependent on their respective charge-to-size ratios [37]. Over the past years, various CZE-MS interfaces, including sheath liquid, nanoflow sheath liquid, and sheathless porous-tip interfaces, have been developed improving overall sensitivity and stability [38]. Still, few works have been reported for native (n)CZE-TDMS in regard to protein analysis [34,35].

Recently, a generic nMS protocol for the analysis of different protein standards (~30 to 230 kDa) on Orbitrap systems was published by our group [13]. The nMS protocol serves as a guide to obtain reproducible and high-quality nMS data and substantiates the novelty and importance of nMS protein analysis. So far, there is no comparable approach reported for nCZE-TDMS, including importance and influence of different method parameters. Herein, we present an operating procedure that utilizes the commercial CZE instrument, SCIEX CESI-8000 Plus, coupled to different Orbitrap instruments for the separation and characterization of native proteins and their complexes. The analyte library covers a wide mass range from ~30 to 800 kDa, including intact-, subunit-, and fragmentation-based analysis. The influence of various CZE method parameters were evaluated, such as BGE composition, acetic acid concentration in the conductive liquid line, and separation voltage. In addition, a universal approach on how to optimize fragmentation settings for metal complex characterization and protein subunit characterization is discussed using selected model analytes. Furthermore, a short section dedicated to general troubleshooting procedures of the nCZE-TDMS setup is included.

## 2 Materials and methods

### 2.1 Materials

A detailed list of all materials used during this study, including vendor and associated product numbers, can be found in **Table S1**.

### 2.2 Preparation of Protein Standards

Protein stock solutions, except for GroEL, were directly desalted and buffer exchanged into 40 mM ammonium acetate (AmAc) using centrifugal filters with 10–50 kDa MWCO, depending on protein molecular weight. First, the filter device was equilibrated with 500 μL of water and spun for 5 min at 12,000×*g*. The protein sample was then dispensed into the filter device, up to a total volume of 500 μL, and spun for 5 min at 12,000×*g* and 4°C or until the sample was concentrated to a volume of 100 μL or less. Ten consecutive buffer exchanges were performed by adding 40 mM AmAc up to 500 μL. Sample was pipetted carefully out of the device and diluted to with 40 mM AmAc to a final concentration of 10 μM. Aliquots were flash frozen and stored at −80°C. In the case of GroEL, a native precipitation was performed before buffer exchange based on a protocol proposed by Freeke *et al.* [39]. Briefly, 1 mg of GroEL was dissolved in 160 μL of buffer 1 (20 mM tris acetate, 50 mM EDTA, 1 mM ATP, 5 mM magnesium chloride) and incubated for 1 h. Subsequently, 40 μL of methanol was added and the solution was incubated for 1 h. Precipitation was performed adding 200 μL of cold acetone (−20 °C) to the sample, vortexed for 30 s, and centrifuged at 21,000×*g* for 10 min at 4°C. Supernatant was carefully removed, and the protein pellet was resuspended in 200 μL of buffer 1. Afterwards, buffer exchange was performed as described above using 100kDa MWCO filters.

### 2.3 Capillary Electrophoresis

A CESI-8000 Plus instrument from SCIEX was used and separation was performed using commercial Neutral OptiMS™ Capillary Cartridges (30 μm inner diameter, 90 cm length) containing an integrated sheathless etched porous nanospray tip. Different BGE compositions ranging from 20 to 60 mM AmAc were used. Before first use, the cartridge was washed and electrically conditioned (see **Table S2** for method details). Each morning, a shortened version of electrical conditioning method was performed applying high voltage for 30 instead of 60 min. For long term storage, the cartridges were rinsed with water and stored at 4 °C.

### 2.4 Mass Spectrometer

The CESI instrument was hyphenated to a custom Q Exactive HF with Extended Mass Range [22] (QE-EMR), a Q Exactive UHMR, and an Orbitrap Eclipse using a Nanospray Flex™ ion source (all from Thermo Fisher Scientific) equipped with an OptiMS Thermo MS Adapter (SCIEX). The spray tip was positioned ~2.5 mm in front of the MS inlet orifice and an ESI voltage between +1.5 and +1.9 kV was applied during separation. To enhance select analyte transmission, in-source collision-induced dissociation (IS-CID) was used in conjunction with Voltage Rollercoaster Filtering (VRF) by optimizing for base peak signal within a user-defined search range [40]. Key MS parameters for MS^1^, MS^2^, and pseudo-MS^3^ experiments for all instruments are listed in **Tables S3-S6**.

### 2.5 Data Analysis

Spectra were analyzed manually using Thermo Xcalibur 4.0 Qual Browser (Thermo Fisher Scientific, Inc.). S/N, taking the highest charge states of a protein distribution in consideration, were calculated as follows: S/N = (NL-B) / (N-B), where NL is the signal intensity, B is the baseline intensity, and N is the noise intensity. Deconvolution of MS^1^ spectra was performed using Unidec GUI version 3.2.0 [41] and detailed parameter settings can be found in **Table S7**. TDValidator v1.0 (Proteinaceous) [42] (max ppm tolerance: 10 ppm; sub ppm tolerance: 5 ppm; cluster tolerance: 0.35; charge range: 1 to at or below that of the analyte; minimum score: 0.5 – 0.73; S/N cutoff: 1 – 5; Distribution Generator: BRAIN; minimum size: 2) was used to assign recorded fragment ions to the primary sequence of analytes. Annotated fragments were manually validated thereafter.

## 3 Results and discussion

### 3.1 Native CZE method development

A standard protein mix composed of carbonic anhydrase II (CA, 29 kDa), alcohol dehydrogenase (ADH, 147 kDa), NIST monoclonal antibody (NIST mAb, 148 kDa), and pyruvate kinase (PK, 231 kDa) was utilized for nCZE-TDMS method development. The mixture covers a wide range of targets including protein-metal complexes, protein-protein complexes, and biopharmaceuticals. Initially, MS^1^ parameters were optimized via direct infusion through the CESI system for each standard individually. A general procedure for this type of experiment can be found at the beginning of the **Supplemental Information**, including a list of MS parameter settings for the QE-EMR instrument (see **Table S3**). The stability of native assemblies is sensitive to changes in MS parameters, especially gas pressures and electric fields applied to the ion optics (e.g. extended trapping, IS-CID). However, the optimal configuration of system parameters can vary strongly depending on the target analyte. For the subsequent nCZE analysis of the native standard mix (nMix), MS settings and concentrations of analytes were adapted accordingly (NIST mAb: 0.5 μM, PK: 1 μM, CA: 3.75 μM, ADH: 2.5 μM). An example electropherogram, using 40 mM AmAc as BGE, a separation voltage of +15 kV and 3 psi supplemental pressure, is depicted in **Figure 1A**. Non-covalent interactions (e.g. tetramer complex) were preserved and typical low charge states were observed for all four analytes as shown in **Figure 1B-G**. A detailed list of all detected species can be found in **Table S8**. Besides the expected main glycoforms for the NIST mAb, mono-glycosylated variants (G0F – G2F) of ~1% relative abundance compared to G0F/G1F could be observed as well (see **Figure 1C-D**). In addition, partial separation of proteoforms containing an additional lysine (+1K) was achieved, which is evident at higher MS resolution (R = 75k at 200 *m/z*, **Figure S1**). ADH, PK, and the singly-truncated form of PK (−21 AA in a single monomer) [43] were detected as tetrameric complexes. In addition, the Zn(II) center of CA was preserved as the main form. Repeatability (n = 5) was established for all four analytes, resulting in RSD values between 0.4 – 1.2% for migration time (MT), 9.1 – 16.5% for peak area, 1.3 – 11.8% for peak intensity, and 10.0 – 15.7% for S/N (see **Table S9**).

**Figure 1.**
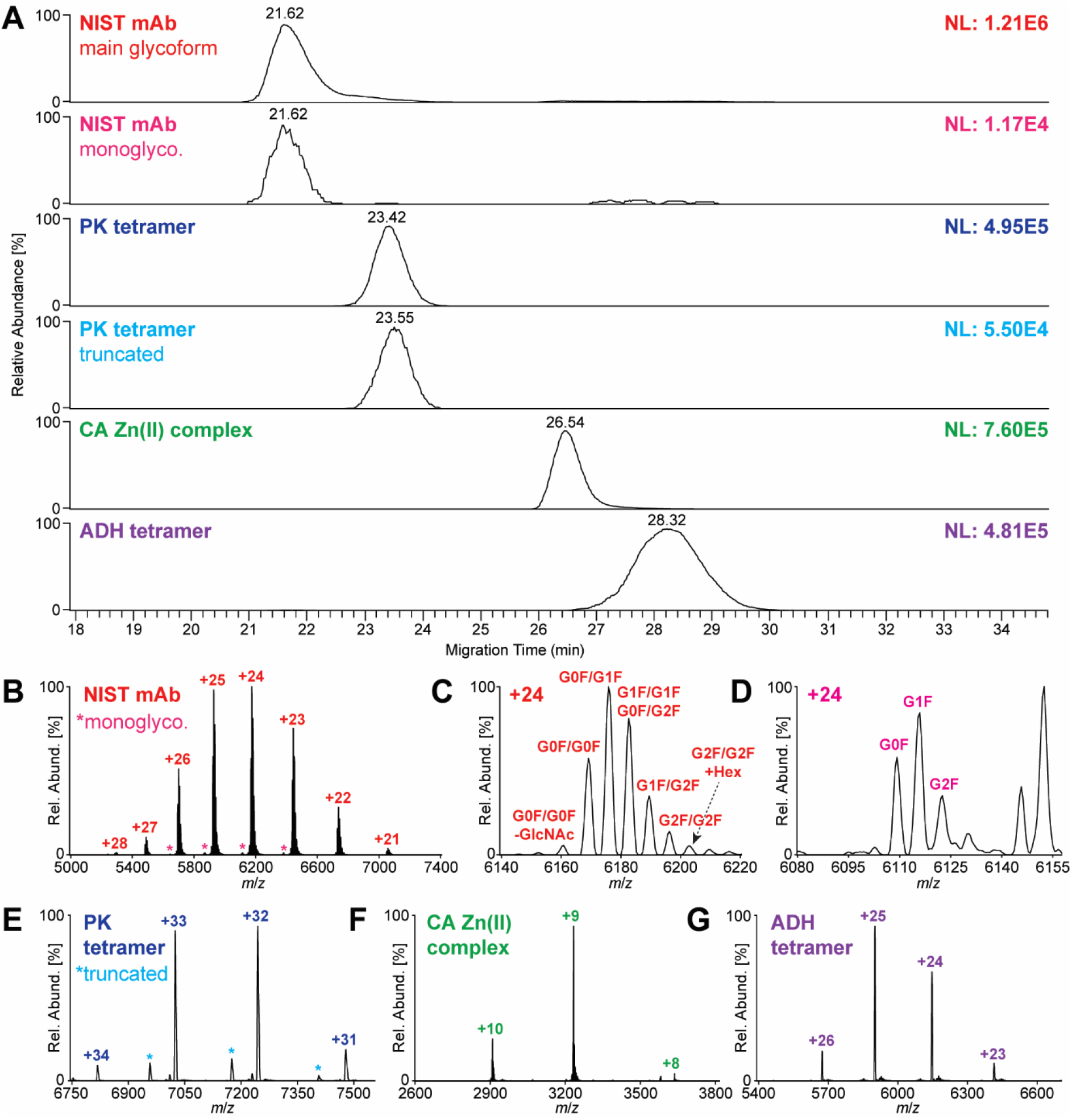
Separation of nMix by nCZE-TDMS using 40 mM AmAc as BGE. (**A**) Extracted Ion Electropherograms (EIEs) of standard protein assemblies created using the three highest charge states, respectively. (**B**) Mass spectrum of NIST mAb. Zoomed in +24 charge state of (**C**) di- and (**D**) monoglycosylated proteoforms of NIST mAb. Mass spectra of (**E**) PK tetramer, (**F**) CA Zn(II) complex, and (**G**) ADH tetramer.

#### 3.1.1 BGE Composition

As a next step, the influence of the concentration of AmAc (20, 40 and 60 mM) in the BGE was investigated (see **Figure 2**). Positioning the sheathless spray tip (2.5 mm from the MS orifice), the voltage required to establish a stable electrospray was affected accordingly: +1.5 kV for 20 mM AmAc, +1.4 kV for 40 mM AmAc, and +1.3 kV for 60 mM AmAc. Alternatively, the spray tip can be moved further away with increasing AmAc concentration. A minimum distance of 2 mm is advised, not only to maintain a stable electrospray but also to avoid contamination of the MS instrument (e.g. by HCl droplets getting sucked in during the preconditioning step of the nCZE method). A low concentration of AmAc (20 mM) affects the separation performance of the nMix negatively; PK overlapped heavily with the NIST mAb signal and lead to peak splitting of the antibody signal. All four species were adequately resolved at 40 mM AmAc despite some moderate NIST mAb tailing. At 60 mM AmAc a similar separation is achieved with marginally narrower peaks and less peak tailing for the NIST mAb signal. However, 60 mM AmAc results in a considerably high current (>5 μA) at +15 kV separation voltage, increasing the risk of damaging the capillary cartridge. For this reason, 40 mM AmAc was utilized for our standard procedure.

**Figure 2.**
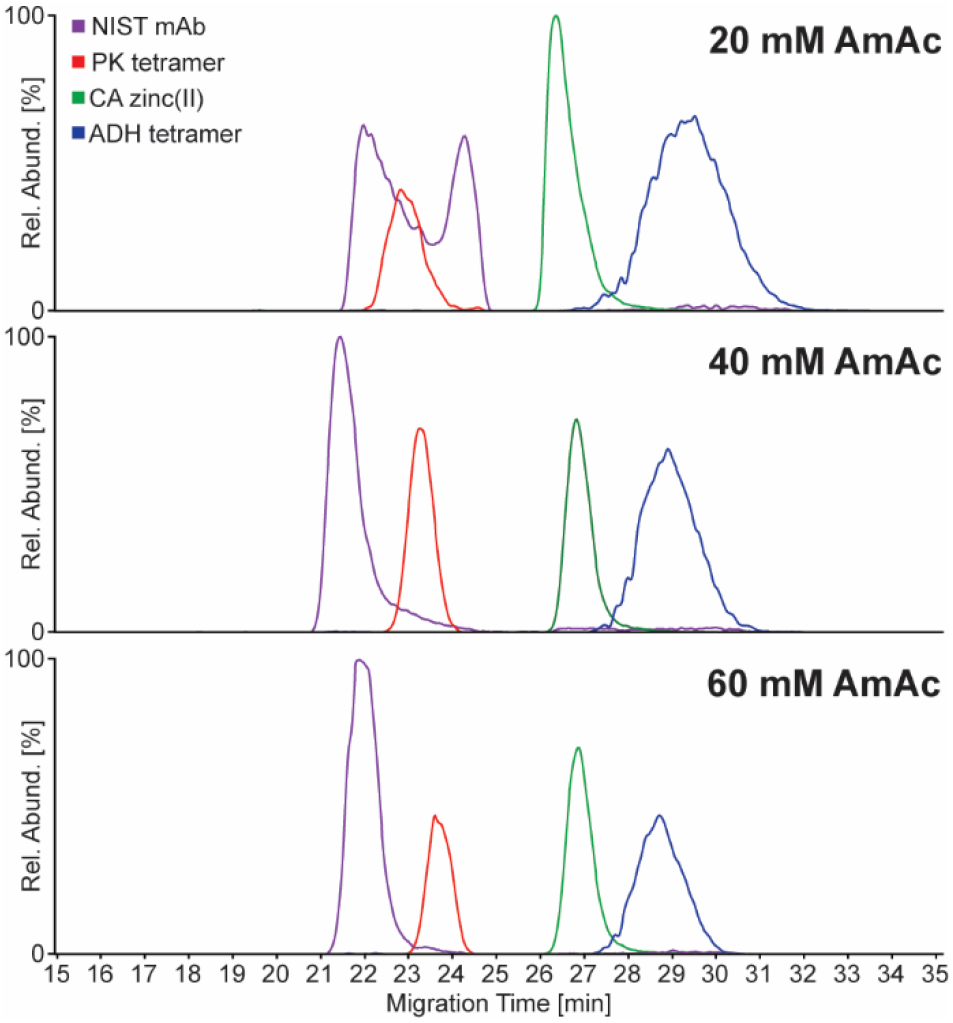
Evaluation of influence of AmAc concentration in BGE on separation performance of nMix by nCZE-TDMS: 20, 40, and 60 mM AmAc. Separation improved with increasing AmAc concentration. EIEs of standard protein assemblies created using the three highest charge states, respectively.

#### 3.1.2 Conductive Liquid (CL) Composition

The membrane-like characteristics of the etched sprayer tip of the CESI-MS adapter is necessary in closing the electrical contact between CL and BGE. The pores are large enough to enable the exchange of protons and small molecules in some cases. Thus, the number of protons delivered by the CL may influence the nature of the native mass spectrum, such as a shift in the observed charged state distribution or a loss of non-covalent interactions due to lowered pH. For this reason, the CL composition was varied between 3 and 10% HAc. In this context, no significant differences were detected (e.g. breakdown of protein complexes), which is in accordance with previous observations [35], and we decided to keep 3% HAc as CL.

#### 3.1.3 Separation Voltage

Besides improving overall separation between analytes, varying the separation voltage can help in understanding the migration behavior of protein assemblies under native conditions. In fact, the estimation of the pI of a protein or protein complex can be quite challenging without detailed knowledge about tertiary/quaternary structures and may differ strongly from what is predicted by the primary sequence of the monomers. As an example, the separation voltage was varied for the analysis of NIST mAb and CA standards (see **Figure S2**). An increase in separation voltage lead to a slightly faster migration of NIST mAb, suggesting a low positive net charge as expected. On the other hand, CA exhibits slower migration in general, and the MT did not change significantly with increasing separation voltage from +10 to +22 kV, indicating a close to zero net charge in solution. For the standard procedure, a separation voltage of +15 kV is recommended.

### 3.2 Optimization of Top-Down MS settings

This section is dedicated to the optimization of TDMS settings for CA, PK, and ADH, including general protocols transferable to other protein targets. Optimization was performed using direct infusion via the CESI system coupled to our Orbitrap Eclipse instrument at a concentration of 2 μM as described in the protocol included in the **Supplemental Information**. MS^1^ parameters for all analytes acquired on the Eclipse instrument can be found in **Table S4**.

In the case of CA we selected the most abundant charge state (+9) using an ion trap isolation width of 100 *m*/*z*. CID was selected as the fragmentation method and applied at 0 normalized collision energy (NCE) to validate the quality of the isolation. Note that collision-, electron-, and ultraviolet-based fragmentation modes are all largely viable and complementary for metalloprotein characterization (see **Figure S3**). The isolation window can be adapted to include or exclude other subspecies (e.g. those with incomplete desolvation) and charge states. After validating the isolation quality, we applied 19 NCE and averaged 100 acquisitions using the conditions described in **Table S5**. Subsequently, we decreased the NCE in increments of 1 down to 14 NCE and repeated the experiment. There are two primary reasons for the repeated experiment across a narrow range. First, different activation energies can yield different fragments which can uniquely contribute to sequence coverage and metal characterization. Second, increases in activation energy can give rise to neutral losses that, in excess, prevent reliable and meaningful interpretation of the data (see **Figure S4**). Thus, the characterization of metalloproteins must strike a fine balance among sufficient metal retention post-activation (low-energy bias), data clarity (low-energy bias), depth of primary sequence coverage (low-energy bias), and total sequence coverage (high-energy bias).

**Figure 3A** shows the fragmentation obtained using 19 NCE of CID and the fragment ions populate the spectrum from ~1,000–5,200 *m*/*z*. TDValidator was used to primarily annotate fragments, and each annotation was manually validated thereafter. As seen in the spectrum, large metal-bound ions (many of which have complementary ions) are present above the *m*/*z* of the isolation window. Many of these annotated fragments, along with their neutral loss distributions, support the presence of zinc (highlighted in blue). As shown in **Figure 3B-E**, this separation of bound and unbound annotations is due to the minimum required size of zinc-containing fragments as opposed to a distinct charge bias imposed by the metal binding.

**Figure 3.**
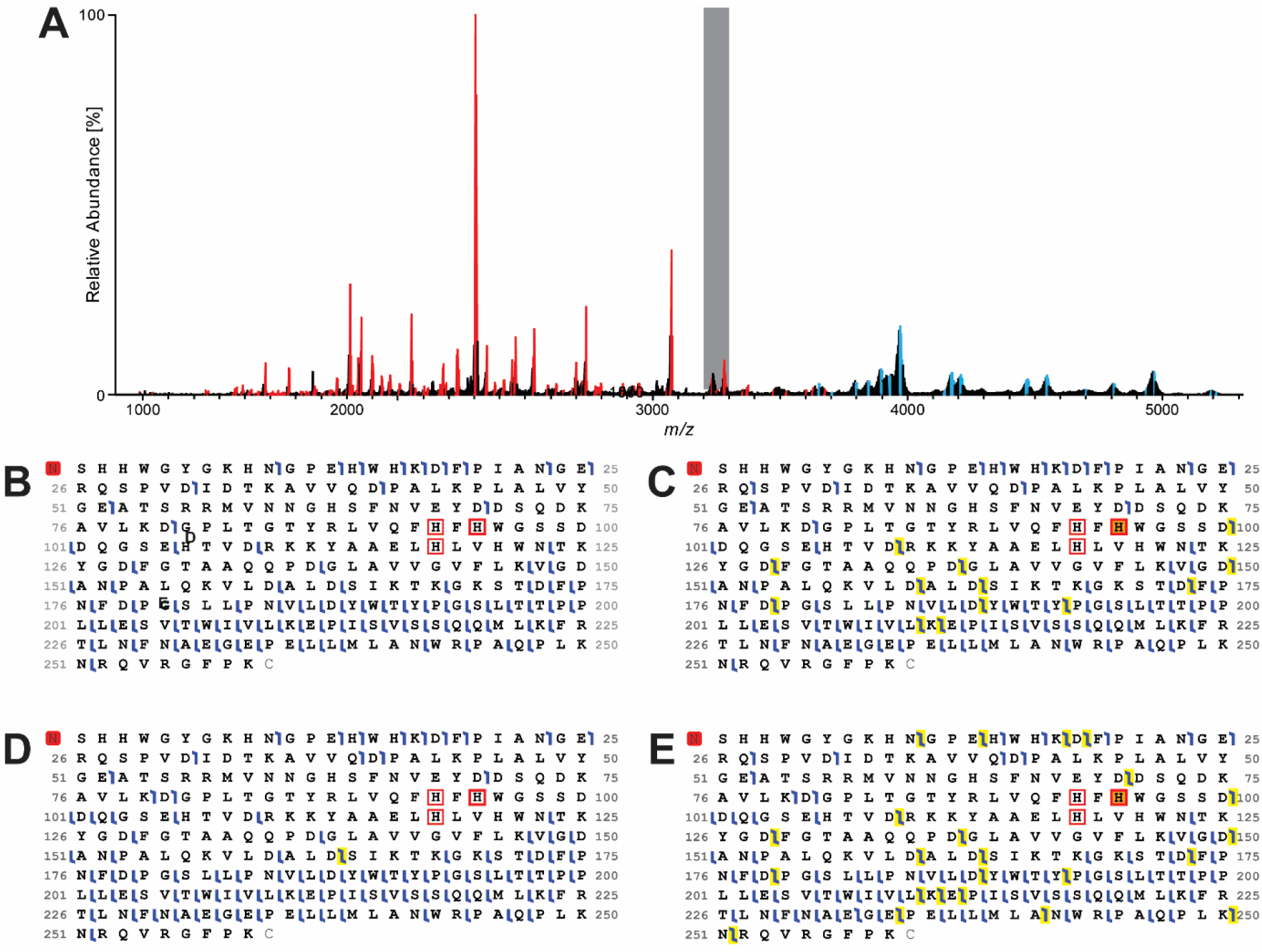
(**A**) CID (19 NCE) Fragmentation spectrum of native CA with highlighted zinc-bound (**blue**) and unbound (**red**) annotated fragments. The gray region denotes the precursor isolation window. (**B-E**) fragmentation maps of CA. Blue flags denote *b*/*y* ions. The red square at the N-terminus indicates an N-terminal acetylation, and the orange square at H95 denotes zinc, with the red square outlines denoting the main participating residues for zinc coordination. (B) CID (19 NCE) fragmentation map of unbound CA. (**C**) CID (19 NCE) fragmentation map of bound CA. The yellow highlights mark the positions of fragments that support the placement of zinc at H95. (**D**) Composite CID (14-19 NCE) fragmentation map of unbound CA. The yellow highlight denotes the one position of the only identified fragment that challenges the placement of zinc at H95. (**E**) Composite CID (14-19 NCE) fragmentation map of bound CA. The yellow highlights mark the positions of fragments that support the placement of zinc at H95.

The importance of preserving non-covalent interactions is amplified when analyzing non-covalent MPCs, including PK as shown in **Figure 4**. Some instruments, such as Orbitrap-Tribrids, possess the ability to select for a precursor at each stage of analysis: subunit ejection and subunit fragmentation. This “true MS3” analysis is not universally available, however. An alternative to true MS3 is “pseudo-MS3,” as depicted in **Figure 4A**. IS-CID is used to eject the monomer without any prior isolation. The monomer is then isolated, in a quadrupole or ion trap, for fragmentation. As this data set was acquired on an Orbitrap Eclipse, it was possible to modify the IS-CID event by employing VRF [40].

**Figure 4.**
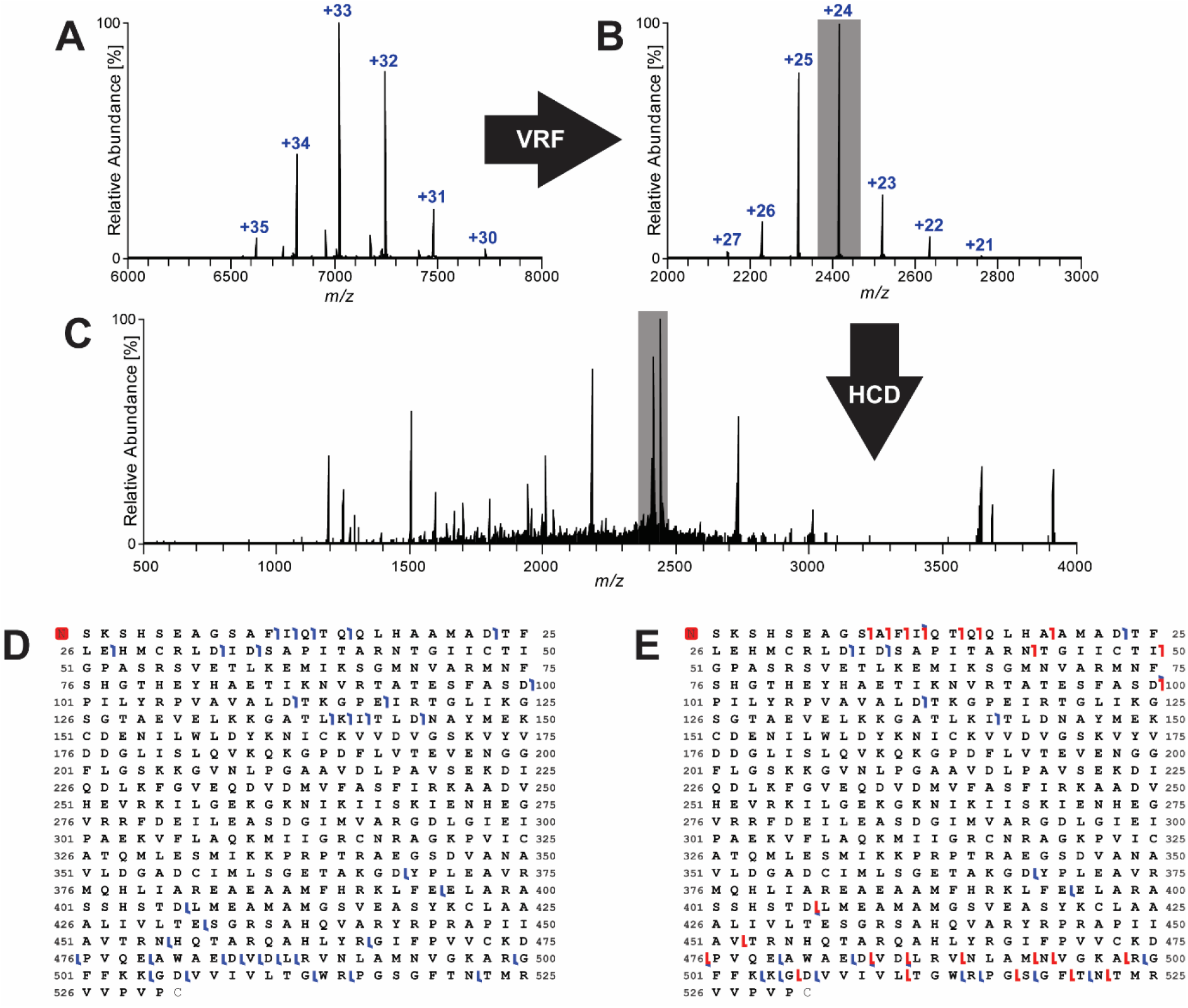
(**A-C**) Activation pathway of PK. The tetramer (**A**) is activated using VRF to yield the monomer (**B**) with optimum transmission. Charge states of the monomer can then be isolated for fragmentation (**C**). (**D-E**) Fragmentation maps of pyruvate kinase summarizing data from (**D**) HCD activation (55-70 V) and (**E**) EThcD activation (1.5 ms ETD reaction time, 50-65 V supplemental HCD activation). The red square at the N-terminus indicates an N-terminal acetylation.

In this method, the voltage profile of the source region is inverted to allow for high activation levels before trading back residual kinetic energy for potential energy for optimal post-activation transmission. VRF can also be used to fragment analytes, as was done for the tetramer (non-covalent dimer) ADH by optimizing for the monomer in a low-resolution mode (see **Figures S5 & 6**). Once ejected, the monomer was fragmented via HCD (**Figure 4D**) and ETD with supplemental HCD activation (**Figure 4E**). The distinct natures of these fragmentation pathways allow for more sequence coverage combined than individually, highlighting the importance of multifaceted characterization.

### 3.3 High molecular weight species

With the recent commercialization of the UHMR Orbitrap instrument, the MS analysis of high molecular weight protein species (>250 kDa) became more accessible. As a showcase, we selected GroEL (~800 kDa), a chaperonin 60 tetradecamer, which has been used as model system for high molecular weight protein complexes in previous studies [44,45]. Analysis was performed using the same CESI settings as for the nMix analysis, with the exception of an increased supplemental pressure (10 instead of 3 psi) as shown in **Figure 5A**. GroEL was detected as intact tetradecamer in accordance with the literature (see **Figure 5B**) [44,45], highlighting the applicability of nCZE for the analysis of high molecular weight protein species.

**Figure 5.**
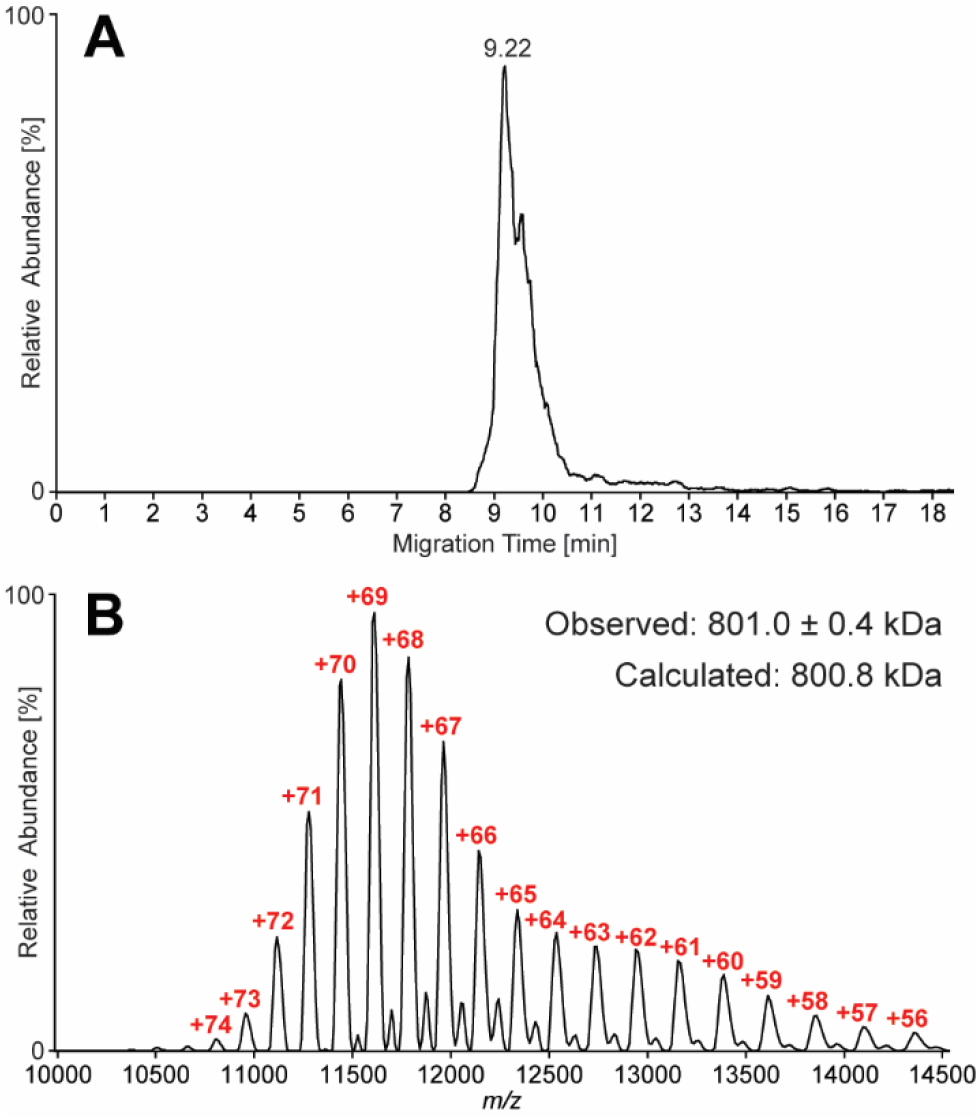
(**A**) Extracted Ion Electropherogram of GroEL standard (3 μM) using the UHMR instrument for detection. (**B**) Mass Spectrum of GroEL confirming presence of main tetradecameric complex ranging from +56 to +74 charge states.

### 3.4 Troubleshooting

Common trouble shooting procedures for the CESI system can be found in the user guide from SCIEX. One of the most common issues encountered by operating the CESI system in native mode is a clogged separation capillary. There are various potential reasons, such as contamination of solutions or precipitation of proteins within the capillary. The latter is more likely to occur during direct infusion experiments using the CESI system if the separation capillary is not immediately purged or placed in the water reservoir after infusion as mentioned previously. If a blockage occurs, as an initial step, it is advisable to resort to the user guide and perform the following actions: attempt to remove the plug by running the washing method (see **Table S2**) while the sprayer tip is submerged in water. If the blockage occurred close to the inlet end of the separation capillary, vacuum can be applied instead. In case that the recommended procedures do not resolve the clogging issue, we came up with a simple but efficient alternative approach. By utilizing the syringe setup depicted in **Figure S7**, it is possible to apply pressure manually, e.g. flushing with water, and remove a potential blockage in the capillary and thus, extend the life span of a supposedly inoperable cartridge. Especially at low flow rates, the AmAc based BGE solution can crystalize at the spray tip, and a stable electrospray cannot be maintained. A video of such an incident was added as **Electronical Supplemental Material**. This phenomenon can occur if the electrospray voltage is set too high or if the spray tip is positioned too close to the MS orifice. A simple solution is to either drop the electrospray voltage or to increase the distance between spray tip and MS orifice. Crystallization may also occur if there is a contaminant at the sprayer tip, e.g. debris or a fiber, which can favor the formation of crystals. Placing the sprayer adapter into the external water reservoir of the CESI system and flush with 0.1 M HCl for 5 min can resolve this issue.

## 4 Concluding remarks

In this work, we focused on establishing and discussing standard procedures for nCZE-TDMS using sheathless ionization analyzing different proteins and protein assemblies. A standard mixture containing ~30 – 230 kDa analytes was established for verifying system performance to promote reproducibility and reliability. Fragmentation parameters were optimized for the native top-down analysis of intact complexes. The demonstrated procedures are considered transferable to a variety of different protein targets. In addition, the capability to analyze high molecular weight protein complexes was demonstrated by analyzing the tetradecamer of GroEL (~800 kDa) using a Q Exactive UHMR instrument for detection. Based on the high-performance separation capability added to nMS, we expect a wide range of potential applications for nCZE-TDMS in the future, including in fields such as structural biology and biomedicine. Additionally, we intend for this study to encourage the CE and MS communities to expand and normalize nCZE-TDMS practices.

## Abbreviations

ADH: alcohol dehydrogenase
AmAc: ammonium acetate
CA: carbonic anhydrase II
CL: conductive liquid
EIEs: Extracted Ion Electropherograms
GELFrEE: Gel-Eluted Liquid Fraction Entrapment Electrophoresis
IS-CID: in-source collision-induced dissociation
*m*/*z*: mass-to-charge ratio
mAb: monoclonal antibody
MPCs: multi-proteoform complexes
MT: migration time
NCE: normalized collision energy
nMix: native standard mix
nMS: native mass spectrometry
nTDMS: native top-down mass spectrometry
PK: pyruvate kinase
QE-EMR: Q Exactive HF with Extended Mass Range
VRF: Voltage Rollercoaster Filtering

## Acknowledgements

This work was supported by the National Institute of General Medical Sciences P41 GM108569 for the National Resource for Translational and Developmental Proteomics at Northwestern University, and the instrumentation award (S10OD025194) from NIH Office of Director. We thank SCIEX for their support including Dr. Fang Wang for the valuable discussions and insightful suggestions throughout this research.

## Conflict of interest

NLK engages in entrepreneurship in the area of top-down proteomics.

