## Supplemental Information 1 for "Standard Procedures for Native CZE-MS of Proteins and Protein Complexes up to 800 kDa"

#### Direct infusion of proteins for $MS^1$ optimization

A general procedure to optimize MS settings for native protein analysis via direct infusion is described in the following:

1) Buffer exchange analyte of interest into the expected BGE solution (model system: 40 mM AmAc).
2) Fill the capillary with protein solution by flushing for 2 min at 100 psi (~2.5 total capillary volumes)
3) Ramp up voltage and pressure over 1 min with similar settings as used for separation (model system: +15 kV, 5 psi)
4) Turn on the Orbitrap instrument and slowly ramp up the electrospray voltage till a spray is established in 100 V increments. Increase to voltage by another 100 to 200 V to ensure stability.
5) Vary MS settings to optimize for S/N (not signal intensity!) of intact protein assembly, ejection efficiency or fragmentation quality depending on objective of experiment.

Additional important remarks: direct infusion experiments of protein solutions via the CESI 8000 plus instrument always bare a certain risk of potentially clogging the capillary due to protein precipitation. Thus, it is advisable to start with low concentrations of proteins (1 µM or lower), especially for higher molecular weight species. In addition, if filled with protein solution but not in operation, the protein solution will start to dry out in the capillary over time starting from the capillary tip. Thus, it is recommended to immediately submerge the capillary end in the water reservoir after the direct infusion is stopped and flush with 0.1 M HCl followed by water for a few minutes to remove remaining protein. Nevertheless, if the separation capillary still gets clogged this way, flushing with 0.1 M HCl for e.g. 10 min usually solves this issue. If that is not the case, please refer to the “troubleshooting” section in the main manuscript.

### Figure S1

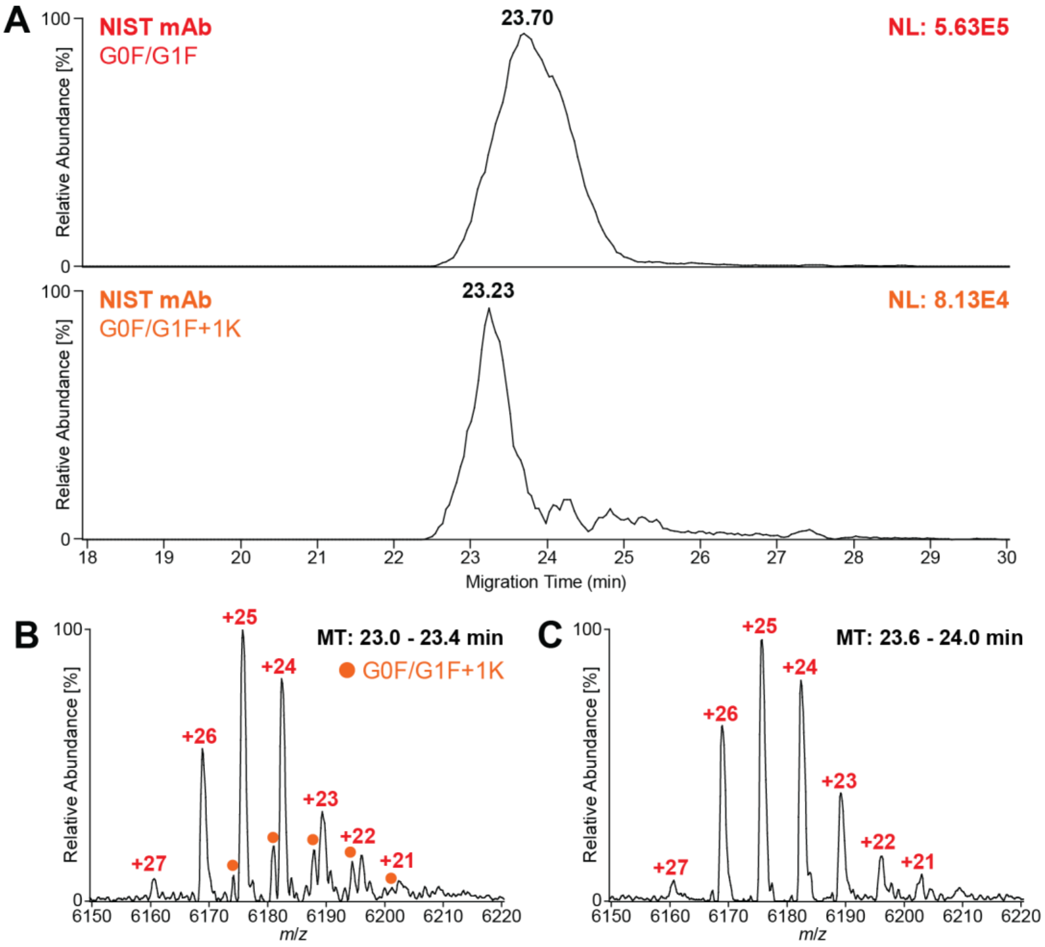


**Figure S1.** Separation of NIST mAb (c = 5 µM) at high MS resolution (R = 75k) using nCZE-TDMS. (**A**) EIEs of main glycoform G0F/G1F (**top**) and G0F/G1F+1K (**bottom**), created using the 3 highest charge states, respectively. (**B**) Mass spectrum generated averaging the MT range from 23.0 – 23.4 min. G0F/G1F+1K related charge states are highlighted (circle, orange). (**C**) Mass spectrum generated averaging the MT range from 23.6 – 24.0 min.

### Figure S2

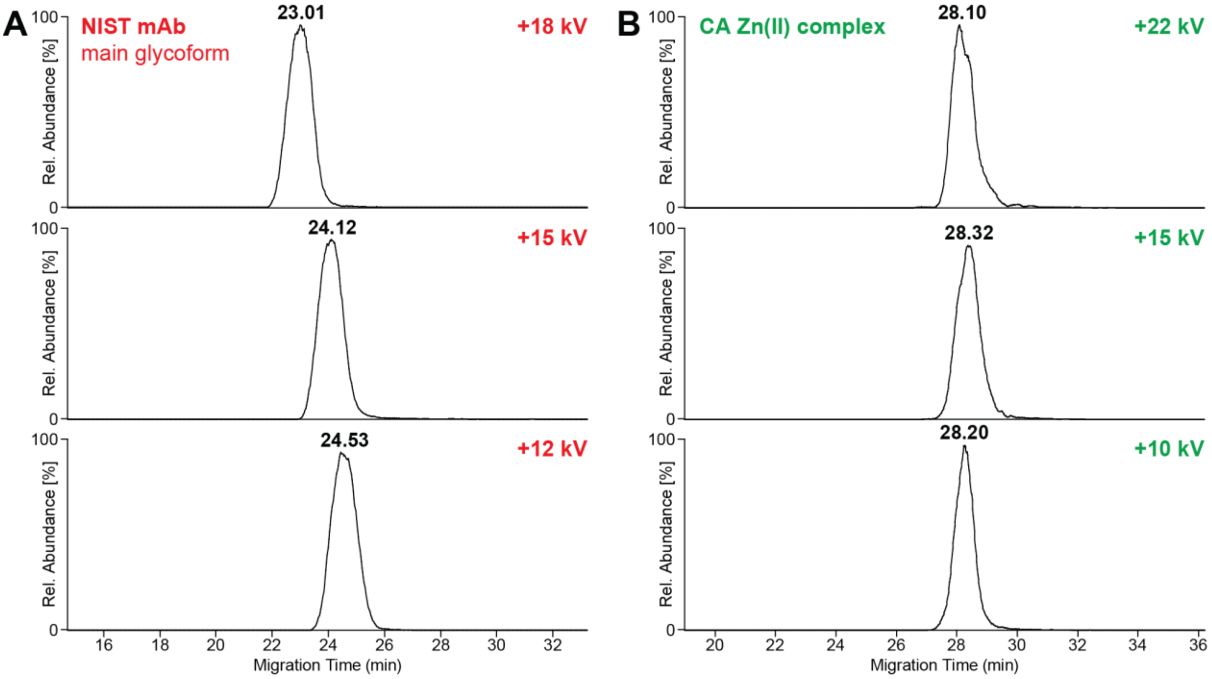


**Figure S2.** Influence of separation voltage on migration behavior of (**A**) NIST mAb and (**B**) CA Zn(II) metal complex using nCZE-TDMS (c = 1 µM, respectively). EIEs of NIST mAb and CA were created using the three highest charge states, respectively. (**A**) Migration time of NIST mAb decreased with increasing voltage, indicating low positive net charge. (**B**) Migration time of CA Zn(II) was not significantly influenced by changes in separation voltage, suggesting a close-to-zero net charge.

### Figure S3

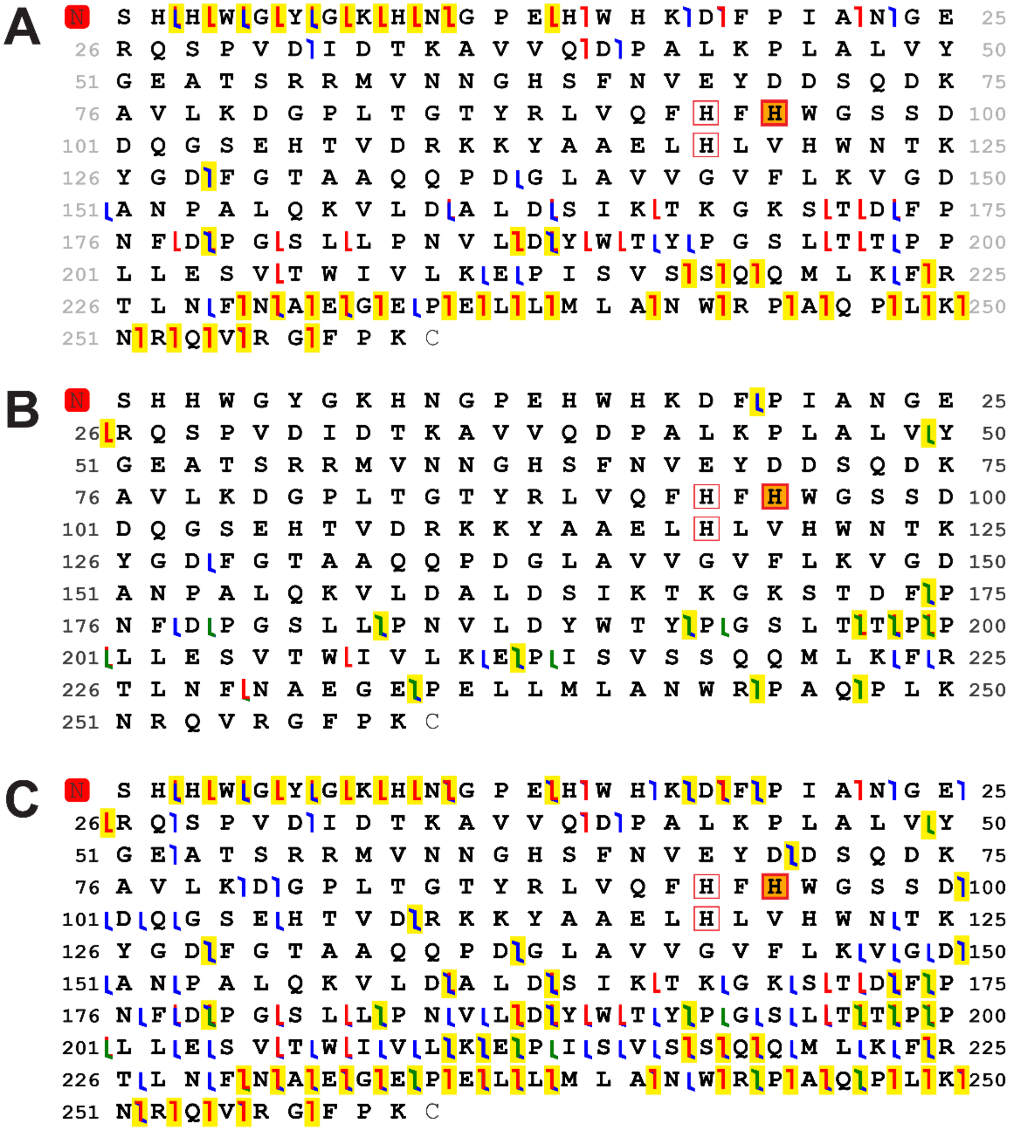


**Figure S3.** Fragmentation maps of CA using (**A**) ETciD (6 ms ETD reaction time, 15 V supplemental CID activation), (**B**) UVPD (17-20 ms activation time), or (**C**) the combined data from all fragmentation types (including the combined CID data from **Figure 3**). Green flags denote *a*/*x* ions, blue flags denote *b*/*y* ions, and red flags denote *c*/*z* ions. The red square at the N-terminus indicates an N-terminal acetylation, and the orange square at H95 denotes zinc, with the red square outlines denoting the main participating residues for zinc coordination. The yellow highlights mark the positions of fragments that support the placement of zinc at H95.

### Figure S4

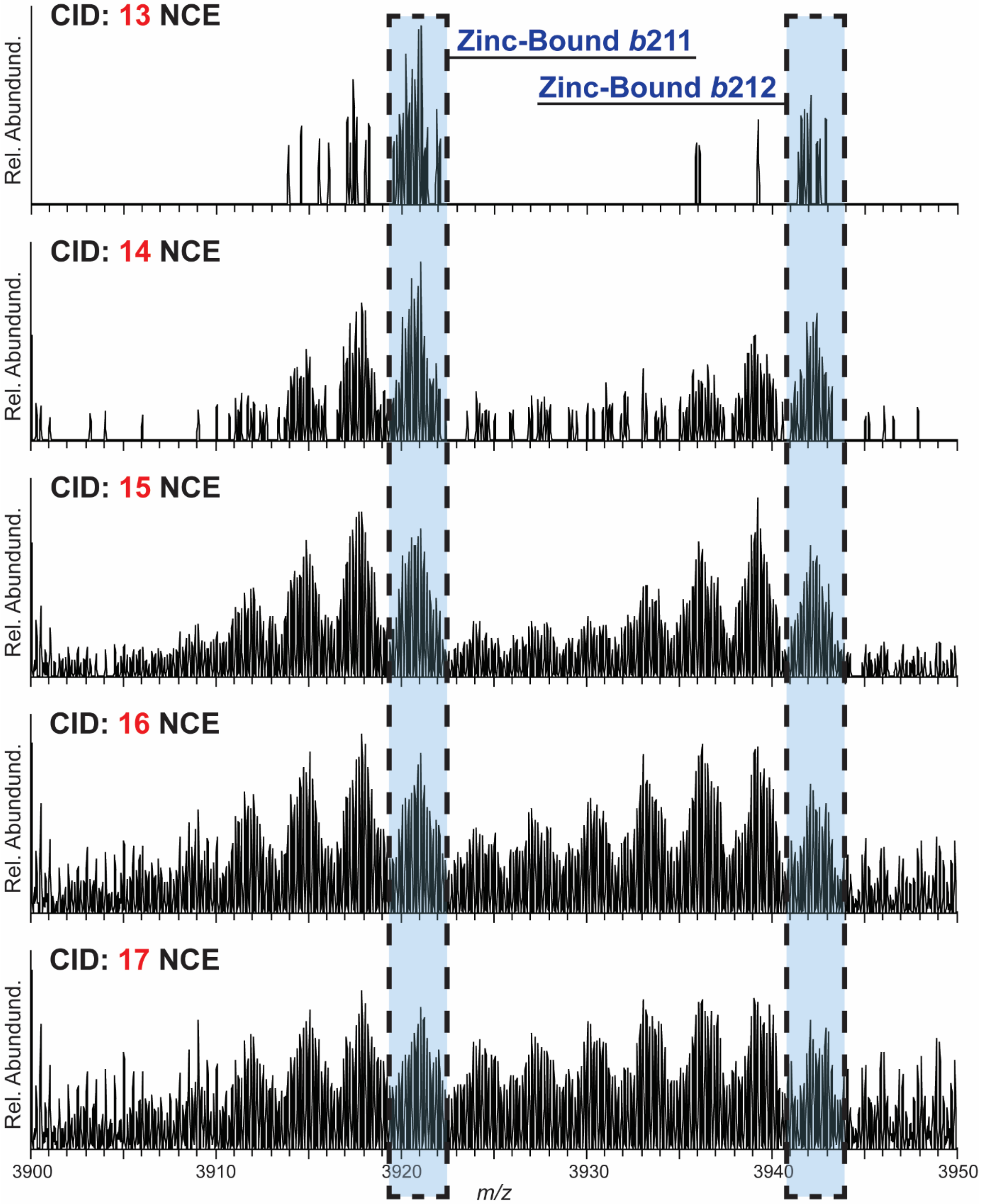


**Figure S4.** Progression of neutral loss distributions with increasing Ion Trap CID activation, leading to a loss of spectral context.

### Figure S5

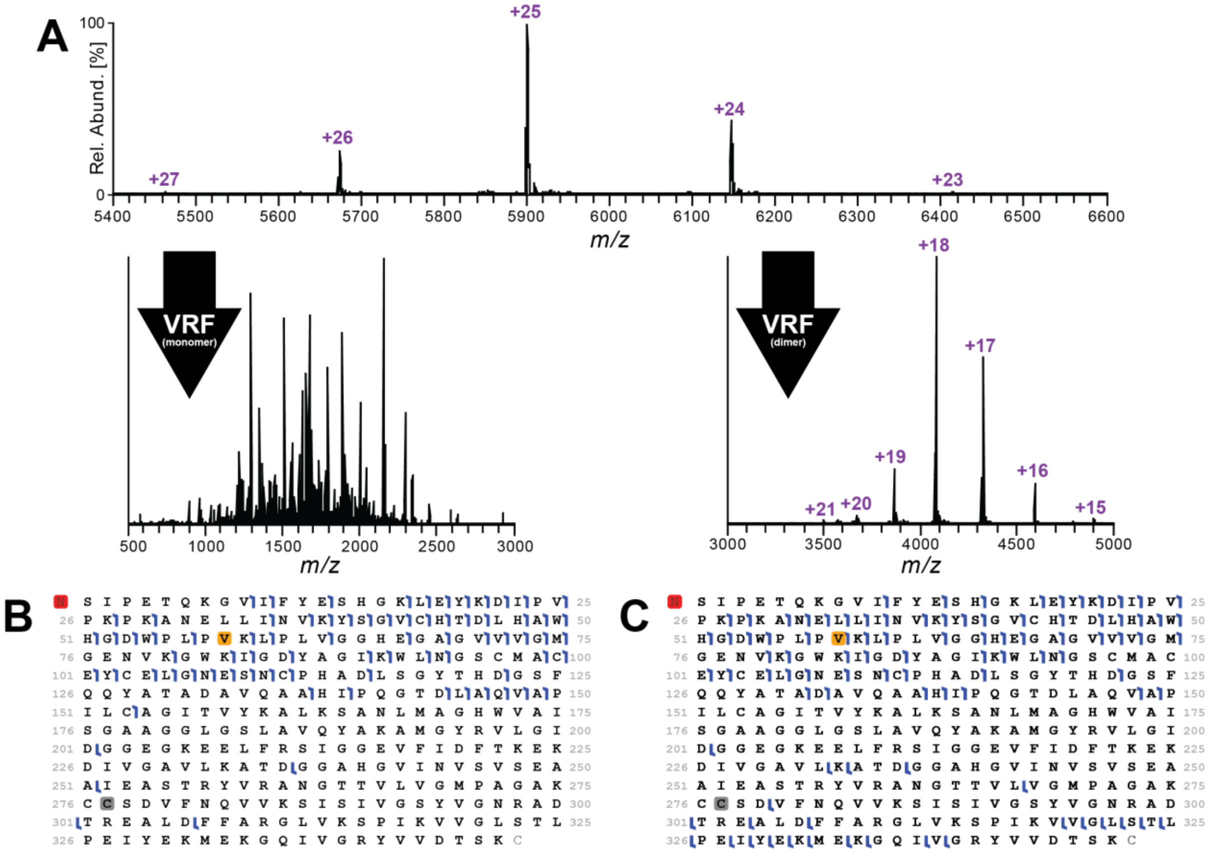


**Figure S5.** (**A**) Activation pathways of ADH. The tetramer (**top**) can be subject to VRF to yield fragments and--at lower resolution--monomers (**left**) or intact dimers (**right**) for further activation. (**B-C**) Fragmentation maps of ADH summarizing data from (B) tailoring VRF for fragmentation and (**C**) activating the ejected dimer (HCD, 35-55 NCE). The red square at the N-terminus indicates an N-terminal acetylation. The yellow square at V58 denotes a V → T sequence variation, and the gray square at C277 denotes a hydrogen loss associated with the formation of a disulfide bridge.

### Figure S6

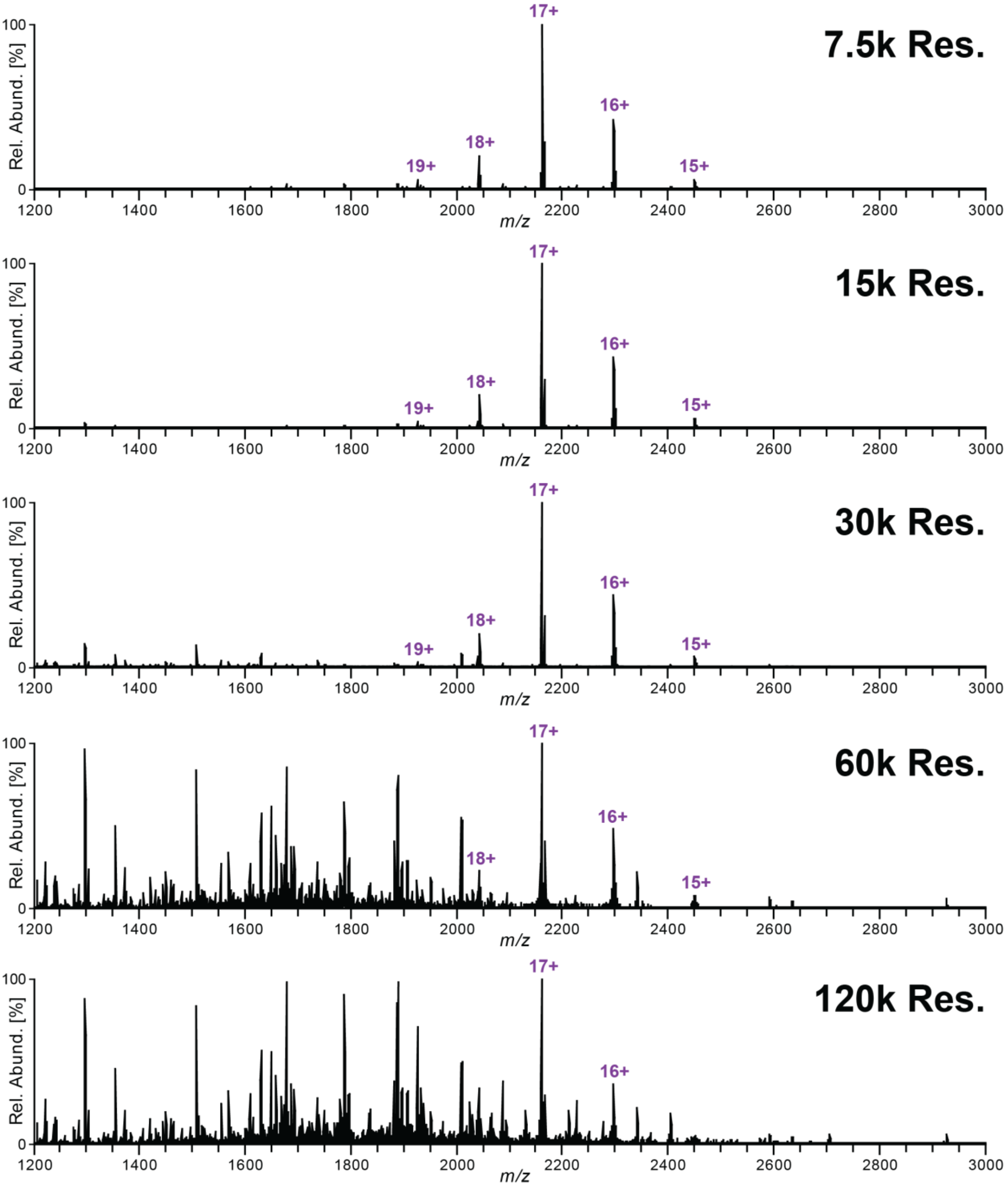


**Figure S6.** Spectra showing VRF-optimized transmission of monomeric ADH under different nominal resolutions, with higher resolutions revealing coincident fragmentation.

### Figure S7

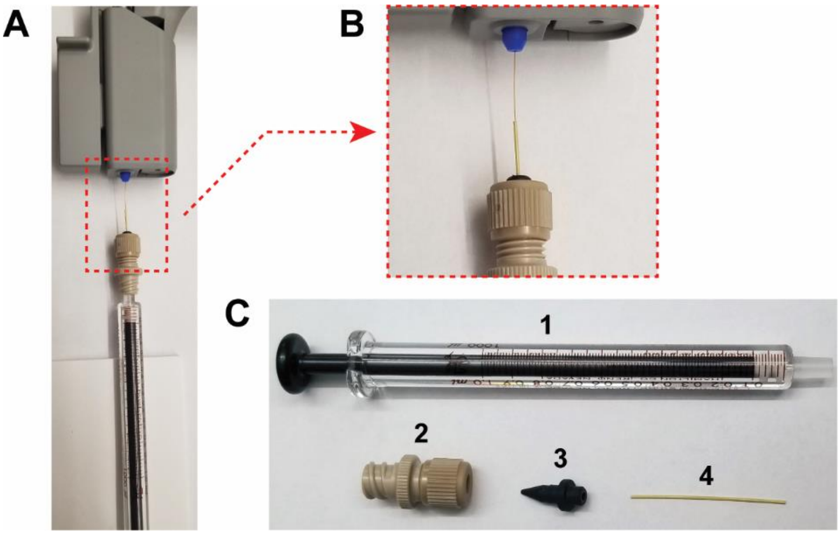


**Figure S7.** (A) Syringe set up for unclogging of separation capillaries. (B) Focus on fitting and connection to capillary inlet. (C) Individual parts: (1), 1 mL syringe, (2) Female Luer-to-MicroTight® Assembly, (3) MicroTight® Ferule, (4) MicroTight® Sleeve .025" OD x .007" ID. An accurate description of vendor and part numbers can be found in the **Table S1**.

**Table S1.** Reagent and Materials List.

| Item | Part Number | Vendor |
|---|---|---|
| **Pyruvate Kinase** from Rabbit Muscle | 10128155001 | Sigma-Aldrich |
| **Alcohol Dehydrogenase** from Baker's Yeast | A8656 | Sigma-Aldrich |
| Bovine **Carbonic Anhydrase** Isozyme II | C2522-5MG | Sigma-Aldrich |
| **NISTmAb**, Humanized IgG1 Monoclonal Antibody | NIST8671 | Sigma-Aldrich |
| Chaperonin 60 (**GroEL**) from Escherichia coli | C7688-1MG | Sigma-Aldrich |
| Ammonium Acetate Solution, for molecular biology, 7.5 M | A2706 | Sigma-Aldrich |
| Acetic Acid, Optima™ LC/MS, Fisher Chemical™ | A11310X1AMP | Fisher Scientific |
| Hyrdrochloric Acid, Certified ACS Plus, 36.5 to 38.0%, Fisher Chemical™ | A144S-500 | Fisher Scientific |
| Water, Optima™ LC/MS Grade, Fisher Chemical™ | W6-4 | Fisher Scientific |
| Methanol, Optima™ LC/MS Grade, Fisher Chemical™ | A456-4 | Fisher Scientific |
| Acetone (Certified ACS), Fisher Chemical™ | A18-4 | Fisher Scientific |
| Tris Acetate-EDTA buffer | T8280 | Sigma-Aldrich |
| Adenosine 5'-Triphosphate (ATP), 10 mM | P0756S | New England Biolabs |
| Magnesium Chloride | M9272-500G | Sigma-Aldrich |
| Amicon Ultra-0.5 mL Centrifugal Filters (10-100 kDa molecular weight cut-off) | UFC503096 | Sigma-Aldrich |
| Nanosray Flex Ion Source | ES260 | Thermo Fisher Scientific |
| CESI Adapter Kit for NanoSpray Flex and Flex NG | B83386 | SCIEX |
| Neutral OptiMS Cartridge | B07368 | SCIEX |
| CESI Vials | B11648 | SCIEX |
| CESI Vial Cap - 100 Pack | B24699 | SCIEX |
| NanoVials - 100 Pack | 5043467 | SCIEX |
| Syringe 1 mL, Model 1001 LT SYR | 81301 | Hamilton |
| MicroTight® Sleeve Yellow .025" OD x .007" ID | F-181 | IDEX |
| Female Luer to MicroTight® Assembly | P-662 | IDEX |

**Table S2.** List of nCZE methods for the CESI 8000 Plus instrument.

| Washing Method | | | | | | |
|---|---|---|---|---|---|---|
| **Time [min]** | **Event** | **Value** | **Duration** | **Inlet** | **Outlet** | **Summary** |
| | Rinse - Pressure | 100 psi | 5.00 min | BI: C1 | BO: A1 | forward |
| | Rinse - Pressure | 100 psi | 10.00 min | BI: A1 | BO: A1 | forward |
| | Rinse - Pressure | 75 psi | 5.00 min | BI: D1 | BO: B1 | reverse |
| | Rinse - Pressure | 100 psi | 30.00 min | BI: D1 | BO: B1 | forward |

| Electrical Conditioning Method | | | | | | |
|---|---|---|---|---|---|---|
| **Time [min]** | **Event** | **Value** | **Duration** | **Inlet** | **Outlet** | **Summary** |
| | Rinse - Pressure | 100 psi | 3.00 min | BI: A1 | BO: A1 | reverse |
| | Rinse - Pressure | 100 psi | 5.00 min | BI: A1 | BO: A1 | forward |
| 0.00 | Separation - Voltage | 15.0 kV | 60.00 min | BI: B1 | BO: A1 | 1.0 min ramp, normal polarity, both |
| 60.00 | Separation - Voltage | 1.0 kV | 5.00 min | BI: B1 | BO: A1 | 5.0 min ramp, normal polarity, both |
| 65.00 | End | | | | | |

| nMix Separation Method | | | | | | |
|---|---|---|---|---|---|---|
| **Time [min]** | **Event** | **Value** | **Duration** | **Inlet** | **Outlet** | **Summary** |
| | Rinse - Pressure | 100 psi | 5.00 min | BI: C1 | BO: A1 | forward |
| | Rinse - Pressure | 100 psi | 3.00 min | BI: A1 | BO: A1 | reverse |
| | Rinse - Pressure | 100 psi | 5.00 min | BI: A1 | BO: A1 | forward |
| | Inject - Pressure | 2.5 psi | 30 sec | SI: A1 | BO: A1 | forward |
| | Wait | | 0.00 min | BI: D1 | BO: A1 | dipping |
| | Inject - Pressure | 2.5 psi | 10 sec | BI: B1 | BO: A1 | forward |
| 0.00 | Separation - Voltage | 15 kV<br>3 psi | 35.00 min | BI: B1 | BO: A1 | 1.0 min ramp, normal polarity, both |
| 1.00 | Relay On | | | | | |
| 35.00 | Separation - Voltage | 1.9 kV<br>5 psi | 5.00 min | BI: B1 | BO: A1 | 5.0 min ramp, normal polarity, both |
| 40.00 | End | | | | | |

BI: A1, B1 = BGE; C1 = 0.1 M HCl, D1 = $H_2O$ SI: A1 = sample BO: A1: CL; B1 = $H_2O$

**Table S3.** Instrument and data acquisition parameters for $MS^1$ analysis using the QE-EMR instrument.

| Parameter | CA | ADH | NIST mAb | PK | nMix |
|---|---|---|---|---|---|
| Acquisition Range [*m/z*] | 500 – 10k | 500 – 10k | 500 – 10k | 500 – 10k | 500 – 10k |
| IS-CID [eV] | off | 30 | 25 | off | 25 |
| Resolution | 15,000 at 200 *m/z* | 15,000 at 200 *m/z* | 15,000 at 200 *m/z* | 15,000 at 200 *m/z* | 15,000 at 200 *m/z* |
| Max Injection Time [ms] | 30 | 30 | 30 | 30 | 30 |
| Number of Microscans | 10 | 10 | 10 | 10 | 10 |
| Extended Trapping [eV] | 50 | 150 | 100 | 150 | 120 |
| HCD Energy [NCE] | off | off | off | off | off |
| Ultra-High Vacuum (UHV) Pressure [mbar]* | $4.16x10^{-10}$ | $6.39x10^{-10}$ | $9.09x10^{-10}$ | $9.09x10^{-10}$ | $7.69x10^{-10}$ |
| Source Temperature [ºC] | 330 | 330 | 330 | 330 | 330 |

*Inner diameter of Peeksil gas restrictor into HCD cell: 100 µm. Adapt HCD pressure to match the pressure values stated.

**Table S4.** Instrument and data acquisition parameters for MS[1] analysis using the Orbitrap Eclipse instrument.

| Parameter | CA | ADH | NIST mAb | PK |
|---|---|---|---|---|
| Acquisition Range [*m*/*z*] | 2k – 8k | 2k – 8k | 2k – 8k | 2k – 8k |
| IS-CID [V] | 15 | 100 | 120 | 140 |
| Source CID compensation scaling | off | -0.12 | -0.15 | -0.18 |
| Resolution | 60,000 at 200 *m/z* | 60,000 at 200 *m/z* | 60,000 at 200 *m/z* | 60,000 at 200 *m/z* |
| Maximum Injection Time [ms] | 100 | 100 | 100 | 100 |
| Number of Microscans | 3 | 3 | 3 | 3 |
| Number of Averaged Acquisitions | 100 | 100 | 100 | 100 |
| HCD Energy [NCE] | off | off | off | off |
| Collision Pressure [mTorr] | 8 | 8 | 8 | 8 |
| Source Temperature [ºC] | 300 | 300 | 300 | 300 |

**Table S5.** Instrument and data acquisition parameters for TDMS analysis using the Orbitrap Eclipse instrument.

| Parameter | CA | ADH* | ADH** | NIST mAb | PK* |
|---|---|---|---|---|---|
| Acquisition Range [*m/z*] | 500 – 6k | 1k – 8k | 500-4k | 1k – 8k | 1k – 8k |
| IS-CID [V] | off | 120 | 220 | 120 | 200 |
| Source CID compensation scaling | off | -0.02 | -0.07 | -0.15 | -0.03 |
| Resolution | 240,000 at 200 m/z | 60,000 at 200 m/z | 60,000 at 200 m/z | 120,000 at 200 m/z | 60,000 at 200 m/z |
| AGC (%) | 100 | 100 | 100 | 100 | 100 |
| Maximum Injection Time [ms] | 100 | 200 | 200 | 350 | 200 |
| Number of Microscans | 3 | 3 | 3 | 3 | 3 |
| Number of Averaged Acquisitions | 100 | 100 | 100 | 100 | 100 |
| HCD Energy [NCE] | off | 35 – 55 | off | 50 | 50 – 70 |
| CID Energy [NCE] | 11 – 19 | off | off | 17 – 50 | 18 – 22 |
| ETD Reaction Time [ms] | 6 | off | off | 2.5 | 1.5 |
| UVPD Activation Time [ms] | 17 – 20 | off | off | 10 – 17 | 8 – 25 |
| Collision Pressure [mTorr] | 8 | 8 | 8 | 8 | 8 |
| Source Temperature [ºC] | 300 | 300 | 300 | 300 | 300 |

*Includes ejection of covalent monomer **Includes direct fragmentation

**Table S6.** Instrument and data acquisition parameters for $MS^1$ analysis of high molecular weight protein assemblies using the UHMR instrument.

| Parameter | GroEl |
|---|---|
| Scan Range [*m/z*] | 1k – 40k |
| IS-CID [eV] | off |
| Resolution | 1,563 at 200 m/z |
| Injection Time [ms] | 50 |
| Number of Microscans | 10 |
| Extended Trapping [eV] | 50 |
| HCD Energy [NCE] | -35 |
| Detector m/z Optimization | High m/z |
| Ion Transfer Target | High *m/z* |
| In-Source Trapping | off |
| Desolvation Voltage [V] | 0 |
| Desolvation Time [ms] | 4 |
| Trapping Voltage [V] | 60 |
| Ultra High Vacuum (UHV) Pressure [mbar]* | 5 ≙ $1.70x10^{-10}$ |
| Source Temperature [ºC] | 310 |

**Table S7.** List of deconvolution parameters for Unidec.

| Decon. Parameter | CA | ADH | NIST mAb | PK |
|---|---|---|---|---|
| *m*/*z* Range [Th] | 2800 - 3700 | 5500 - 6750 | 5150 - 7350 | 6700 - 7600 |
| Bin Every | 0 | 0 | 0 | 0 |
| Charge Range | 7 - 10 | 23 - 26 | 20 - 28 | 30 - 35 |
| Mass Range [Da] | 28k - 30k | 145k – 150k | 145k – 150k | 227k – 235k |
| Sample Mass Every [Da] | 0.1 | 0.1 | 0.1 | 0.1 |
| Peak Detection Range [Da] | 5 | 5 | 5 | 5 |
| Peak Detection Threshold | 0.01 | 0.5 | 0.002 | 0.02 |

**Table S8.** List of protein species observed analyzing the model protein mix (n = 5) by nCZE-TDMS with the QE-EMR as detector. The most abundant proteoforms are highlighted (**bold**).

| Protein | Species | Theoretical Average Mass (Da) | Observed Average Mass (Da) | Mass Error (Da) | Mass Error (ppm) |
|---|---|---|---|---|---|
| NIST | G0F | 146,592.5 | 146,593.6 ±1.9 | 1.2 | 8.0 |
| | G1F | 146,754.6 | 146,754.1 ±2.3 | -0.5 | -3.7 |
| | G2F | 146,916.8 | 146,913.1 ±1.6 | -3.6 | -24.8 |
| | G0F/G0F-GlcNac | 147,834.6 | 147,833.0 ±0.6 | -1.6 | -10.8 |
| | G0F/G0F | 148,037.8 | 148,038.1 ±1.0 | 0.3 | 1.9 |
| | **G0F/G1F** | **148,200.0** | **148,201.9 ±0.5** | **1.9** | **13.0** |
| | G1F/G1F, G0F/G2F | 148,362.1 | 148,361.9 ±1.3 | -0.2 | -1.1 |
| | G1F/G2F | 148,524.3 | 148,523.5 ±1.3 | -0.8 | -5.3 |
| | G2F/G2F | 148,686.4 | 148,685.1 ±1.8 | -1.3 | -8.5 |
| | G2F/G2F+Hex | 148,848.5 | 148,843.1 ±0.3 | -5.4 | -36.3 |
| | G2F/G2F+2Hex | 149,010.7 | 149,006.3 ±1.5 | -4.3 | -29.2 |
| PK | Truncated | 229,564.2 | 229,563.2 ±1.3 | -1.0 | -4.4 |
| | **Tetramer** | **231,775.6** | **231,778.2 ±0.7** | **2.6** | **11.2** |
| CA | Loss $H_2O$+Ac | 29,027.9 | 29,028.0 ±0.5 | 0.1 | 4.4 |
| | Loss Ac | 29,045.9 | 29,045.1 ±0.7 | -0.8 | -27.1 |
| | Loss H2O | 29,069.9 | 29,069.7 ±0.0 | -0.2 | -6.4 |
| | **Zn(II) complex** | **29,087.9** | **29,088.1 ±0.0** | **0.2** | **6.9** |
| | $Na^+$ adduct | 29,109.9 | 29,109.5 ±0.1 | -0.4 | -12.4 |
| | $K^+$ adduct | 29,126.0 | 29,127.1 ±0.7 | 1.1 | 38.2 |
| | $HCO_3^-$ adduct | 29,148.9 | 29,148.8 ±0.5 | -0.1 | -4.6 |
| ADH | **Tetramer** | **147,492.1** | **147,495.0 ±1.0** | **2.9** | **19.9** |

GlcNAc = N-Acetylglucosamine, Hex = Hexose

**Table S9.** Repeatability of nMix analysis using nCZE-TDMS using the most abundant proteoforms, respectively.

| Analyte | Run # | MT [min] | Peak Area | Peak Intensity | S/N |
|---|---|---|---|---|---|
| NIST (G0F/G1F) | 1 | 21.56 | 5.10E07 | 1.25E06 | 944 |
| | 2 | 21.69 | 6.49E07 | 1.21E06 | 1231 |
| | 3 | 21.56 | 6.34E07 | 1.25E06 | 1173 |
| | 4 | 21.35 | 5.91E07 | 1.24E06 | 880 |
| | 5 | 21.22 | 6.08E07 | 1.22E06 | 913 |
| | Mean | 21.48 | 5.99E07 | 1.23E06 | 1028 |
| | StDev | 0.19 | 5.42E06 | 1.59E04 | 162 |
| | RSD [%] | 0.88 | 9.05 | 1.29 | 15.72 |
| PK | 1 | 23.47 | 1.59E07 | 4.63E05 | 470 |
| | 2 | 23.36 | 1.86E07 | 4.91E05 | 558 |
| | 3 | 23.44 | 2.20E07 | 5.52E05 | 615 |
| | 4 | 23.42 | 2.18E07 | 5.48E05 | 600 |
| | 5 | 22.89 | 2.08E07 | 5.41E05 | 591 |
| | Mean | 23.32 | 1.98E07 | 5.19E05 | 567 |
| | StDev | 0.24 | 2.57E06 | 3.98E04 | 58 |
| | RSD [%] | 1.04 | 13.00 | 7.67 | 10.23 |
| CA | 1 | 26.60 | 3.62E07 | 1.16E06 | 1285 |
| | 2 | 26.54 | 3.47E07 | 1.16E06 | 1251 |
| | 3 | 26.52 | 4.21E07 | 1.36E06 | 1445 |
| | 4 | 26.49 | 3.90E07 | 1.34E06 | 1412 |
| | 5 | 26.73 | 5.16E07 | 1.02E06 | 1605 |
| | Mean | 26.58 | 4.07E07 | 1.21E06 | 1399 |
| | StDev | 0.10 | 6.72E06 | 1.43E05 | 141 |
| | RSD [%] | 0.36 | 16.50 | 11.81 | 10.08 |
| ADH | 1 | 28.45 | 5.70E07 | 6.61E05 | 1319 |
| | 2 | 28.40 | 5.08E07 | 5.65E05 | 1187 |
| | 3 | 28.40 | 5.69E07 | 6.45E05 | 1330 |
| | 4 | 28.13 | 6.01E07 | 7.12E05 | 1381 |
| | 5 | 29.06 | 7.19E07 | 7.39E05 | 1563 |
| | Mean | 28.49 | 5.93E07 | 6.64E05 | 1356 |
| | StDev | 0.34 | 7.77E06 | 6.71E04 | 136 |
| | RSD [%] | 1.21 | 13.10 | 10.10 | 10.02 |

Stdev = standard deviation, RSD = relative standard deviation
